# Software for the integration of multi-omics experiments in Bioconductor

**DOI:** 10.1101/144774

**Authors:** Marcel Ramos, Lucas Schiffer, Angela Re, Rimsha Azhar, Azfar Basunia, Carmen Rodriguez Cabrera, Tiffany Chan, Philip Chapman, Sean Davis, David Gomez-Cabrero, Aedin C. Culhane, Benjamin Haibe-Kains, Kasper D. Hansen, Hanish Kodali, Marie Stephie Louis, Arvind Singh Mer, Markus Riester, Martin Morgan, Vincent Carey, Levi Waldron

## Abstract

Multi-omics experiments are increasingly commonplace in biomedical research, and add layers of complexity to experimental design, data integration, and analysis. R and Bioconductor provide a generic framework for statistical analysis and visualization, as well as specialized data classes for a variety of high-throughput data types, but methods are lacking for integrative analysis of multi-omics experiments. The MultiAssayExperiment software package, implemented in R and leveraging Bioconductor software and design principles, provides for the coordinated representation of, storage of, and operation on multiple diverse genomics data. We provide all of the multiple ‘omics data for each cancer tissue in The Cancer Genome Atlas (TCGA) as ready-to-analyze MultiAssayExperiment objects, and demonstrate in these and other datasets how the software simplifies data representation, statistical analysis, and visualization. The MultiAssayExperiment Bioconductor package reduces major obstacles to efficient, scalable and reproducible statistical analysis of multi-omics data and enhances data science applications of multiple omics datasets.

## INTRODUCTION

Multi-assay experiments collect multiple, complementary data types for a set of specimens. Bioconductor provides classes to ensure coherence between a single assay and patient data during data analysis, such as eSet and SummarizedExperiment. However, novel challenges arise in data representation, management, and analysis of multi-assay experiments (4), that cannot be addressed by these or other single-assay data architectures. These include 1) coordination of different assays on, for example, genes, microRNAs, or genomic ranges, 2) coordination of missing or replicated assays, 3) sample identifiers that differ between assays, 4) re-shaping data to fit the variety of existing statistical and visualization packages, 5) doing the above in a concise and reproducible way that is amenable to new assay types and data classes.

The need for a unified data model for multi-omics experiments has been recognized in other projects, such as MultiDataSet (9) and CNAMet (10). Our developments are motivated by an interest in bridging effective single-assay API elements, including endomorphic feature and sample subset operations, to multi-omic contexts of arbitrary complexity and volume (**Supplemental Table 1**). A main concern in our work is to allow data analysts and developers to simplify the management of both traditional in-memory assay stores for smaller datasets, and out-of-memory assay stores for very large data, in such formats as HDF5, tabix-indexed VCF, or Google BigTable.

MultiAssayExperiment provides data structures and methods for representing, manipulating, and integrating multi-assay genomic experiments. It integrates an *open-ended* set of R and Bioconductor single-assay data classes, while abstracting the complexity of back-end data objects a sufficient set of data manipulation, extraction, and re-shaping methods to interface with most R/Bioconductor data analysis and visualization tools. We demonstrate its use by representing unrestricted data from TCGA as a single MultiAssayExperiment object per cancer type, and demonstrating greatly simplified multi-assay analyses with these and other public multi-omics datasets.

## MATERIALS AND METHODS

MultiAssayExperiment <u>https://bioconductor.org/packages/**MultiAssayExperiment**</u> introduces a Bioconductor S4 class with three key components: 1) *colData*, a “primary” dataset containing patient or cell line-level characteristics such as pathology and histology, 2) *ExperimentList*, a list of results from complementary experiments, and 3) *sampleMap*, a map that relates these elements (Figure 1a). *ExperimentList* data elements may be of any data class that has standard methods for basic subsetting (single square bracket `[`) and dimension names and sizes (`dimnames()` and `dim()`). Key methods for the MultiAssayExperiment class include:

1. A constructor function and associated validity checks that simplifies creating MultiAssayExperiment objects, while allowing for flexibility in representing complex experiments.
2. Subsetting operations allowing data selection by genomic identifiers or ranges, clinical/pathological variables, available complete data (subsets that include no missing values), and by specific assays.
3. Robust and intuitive extraction and replacement operations for components of the *MultiAssayExperiment*.

**Figure 1.**
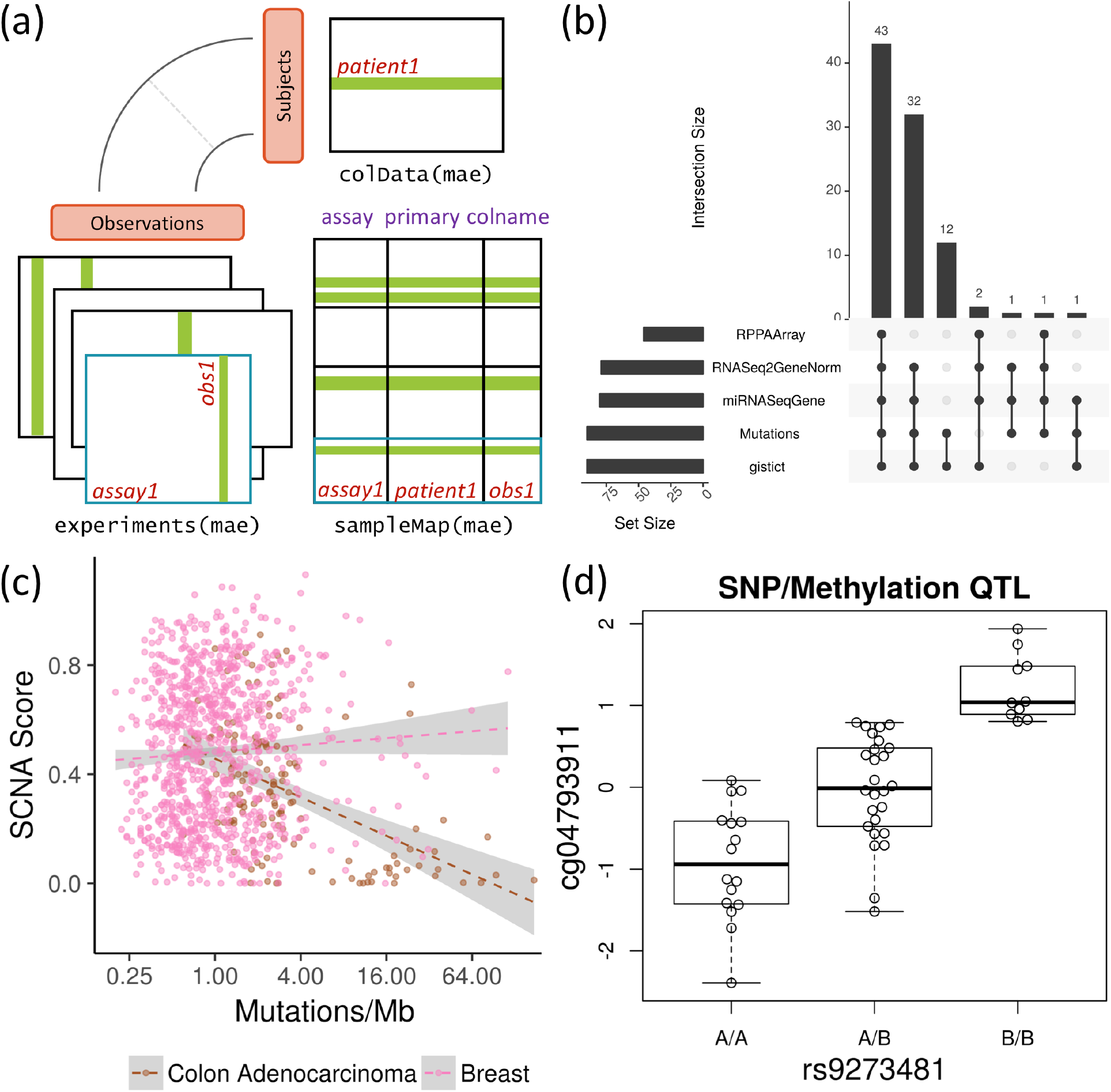
MultiAssayExperiment design and applications. **(a) The MultiAssayExperiment object schematic** shows the design of the infrastructure class. The colData provides data about the patients, cell lines, or other biological units, with one row per unit and one column per variable. The experiments are a list of assay datasets of arbitrary class, with one column per observation. The sampleMap links a single table of patient data (colData) to a list of experiments via a simple but powerful table of experiment:patient edges (relationships), that can be created automatically in simple cases or in a spreadsheet if assay-specific sample identifiers are used. sampleMap relates each column (observation) in the assays (experiments) to exactly one row (biological unit) in colData; however, one row of colData may map to zero, one, or more columns per assay, allowing for missing and replicate assays. Green stripes indicate a mapping of one subject to multiple observations across experiments. **(b) The UpSetR graphic** represents a complex Venn diagram of assay availability for patients in a MultiAssayExperiment. This Glioblastoma object has been subset to only four of its original 12 assays. The barplot on the left shows sample size of each experiment; links to its right indicate combinations of 1 to 4 experiments, with bars above showing the number of patients having exactly those data types. **(c) Extent of copy number alteration vs. somatic mutation burden**. Cancer types with high levels of aneuploidy often show a positive correlation of mutation load and chromosomal instability (CIN) (12), perhaps due to a higher tolerance of deleterious mutations, as shown here in orange for breast cancer. Tumors with a hypermutator phenotype rarely display extensive CIN, resulting in a negative correlation of mutation load and CIN in cancer types where hypermutation is common (shown in grey for colon adenocarcinoma). **(d) Methylation Quantitative Trait Locus** identified from an on-disk representation of Variant Call Format (VCF) files of the 1000 Genomes Project integrated with 450K methylation array data as a MultiAssayExperiment.

The *MultiAssayExperiment* API is based wherever possible on *SummarizedExperiment* while supporting heterogeneous multi-omics experiments. MultiAssayExperiment design, constructor, subsetting, extraction, and helper methods, as well as methods and code for the examples demonstrated here, are detailed in the **Supplemental Methods**.

## RESULTS

The MultiAssayExperiment class and methods (Table 1) provide a flexible framework for integrating and analyzing complementary assays on an overlapping set of samples. It integrates any data class that supports basic subsetting and dimension names, so that many data classes are supported by default without additional accommodations. The MultiAssayExperiment class (Figure 1a) ensures correct alignment of assays and patients, provides coordinated subsetting of samples and features while maintaining correct alignment, and enables simple integration of data types to formats amenable to analysis by existing tools. Basic usage is outlined in Video 1 (https://youtu.be/w6HWAHaDpyk), and in the *QuickStartMultiAssay* vignette accompanying the package.

**Table 1.** Summary of the MultiAssayExperiment API.

| Category and Function | Description | Returned class |
| --- | --- | --- |
| **Constructors** |  |  |
| MultiAssayExperiment | Create a MultiAssayExperiment object | MultiAssayExperiment |
| ExperimentList | Create an ExperimentList from a List or list | ExperimentList |
| **Accessors** |  |  |
| colData | Get or set data that describe the samples | DataFrame |
| experiments | Get or set the list of experimental data objects as original classes | ExperimentList |
| assays | Get the list of experimental data numeric matrices | SimpleList |
| assay | Get the first experimental data numeric matrix | matrix, matrix-like |
| sampleMap | Get or set the map relating observations to subjects | DataFrame |
| metadata | Get or set additional data descriptions | list |
| rownames | Get row names for all experiments | CharacterList |
| colnames | Get column names for all experiments | CharacterList |
| **Subsetting** |  |  |
| mae[ i, j, k ] | Get rows, columns, and/or experiments | MultiAssayExperiment |
| mae[ i, , ] | GRanges, character, integer, logical, List, list | MultiAssayExperiment |
| mae[ , j, ] | character, integer, logical, List, list | MultiAssayExperiment |
| mae[ , , k ] | character, integer, logical | MultiAssayExperiment |
| mae[[ i ]] | Get or set object of arbitrary class from experiments | (varies) |
| mae[[ i ]] | character, integer, logical |  |
| mae\$column | Get or set colData column | vector (varies) |
| **Management** |  |  |
| complete.cases | Identify subjects with complete data in all experiments | vector (logical) |
| duplicated | Identify subjects with replicate observations per experiment | list of LogicalLists |
| mergeReplicates | Merge replicate observations within each experiment, using function | MultiAssayExperiment |
| intersectRows | Return features that are present for all experiments | MultiAssayExperiment |
| intersectColumns | Return subjects with data available for all experiments | MultiAssayExperiment |
| prepMultiAssay | Troubleshoot common problems when constructing main class | list |
| **Reshaping** |  |  |
| longFormat | Return a long and tidy DataFrame with optional colData columns | DataFrame |
| wideFormat | Create a wide DataFrame, 1 row per subject | DataFrame |
| **Combining** |  |  |
| c | Concatenate an experiment | MultiAssayExperiment |

Note. assay refers to a procedure for measuring the biochemical or immunological activity of a sample, e.g. RNA-seq, segmented copy number, and somatic mutation calls would be considered three different assays. experiment refers to the application of an assay to a set of samples. In general it is assumed that each experiment uses a different assay type, although an assay type may of course be repeated in different experiments. mae refers to a MultiAssayExperiment object. subject refers to patient, cell line, or other biological unit. observation refers to results of an assay, e.g. gene expression, somatic mutations, etc. features refer to measurements returned by the assays, labeled by row names or genomic ranges.

We coordinated over 300 assays on 19,000 specimens from 33 different cancers in The Cancer Genome Atlas, along with curated clinical data and published subtypes, and represented these as one MultiAssayExperiment per cancer type (**Supplemental Table 3**). These data objects link each assay to their patient of origin, allowing more straightforward selection of cases with complete data for assays of interest, and integration of data across assays and between assays and clinical data. We demonstrate applications of MultiAssayExperiment for visualizing the overlap in assays performed for adrenocortical carcinoma patients (Figure 1b), confirming recently reported correlations between somatic mutation and copy number burden in colorectal cancer and breast cancer (Figure 1c), identifying a SNP/methylation quantitative trait locus using remotely stored tabix-indexed VCF files for the 1000 genomes project (Figure 1d), calculating correlations between copy number, gene expression, and protein expression in the NCI-60 cell lines (**Supplemental Figure 3**), and multi-assay gene set analysis for ovarian cancer (**Supplemental Figures 1-2**). Demonstrative code chunks and fully reproducible scripts are given to demonstrate the simple and powerful flexibility provided by MultiAssayExperiment.

## DISCUSSION

MultiAssayExperiment enables coordinated management management and extraction of complex multi-assay experiments and clinical data, with the same ease of user-level coding as for a single experiment. Its extensible design supports any assay data class meeting basic requirements, including out-of-memory representations for very large datasets. We have confirmed “out-of-the-box” compatibility with on-disk data representations, including the DelayedMatrix class via an HDF5 backend (11), and the VcfStack class based on the GenomicFiles infrastructure. Future work will focus on higher-level visualization, integration, and analysis tools using MultiAssayExperiment as a building block. This project will receive long-term support as a necessary element of multi-assay data representation and analysis in Bioconductor.

## ACKNOWLEDGEMENTS

The authors’ work was funded by the National Cancer Institute of the National Institutes of Health [U24CA180996 to MM]. This work was supported by the CUNY High Performance Computing Center, which is operated by the College of Staten Island and funded, in part, by grants from the City of New York, State of New York, CUNY Research Foundation, and National Science Foundation Grants CNS-0958379, CNS-0855217 and ACI 1126113.

